# *In vivo* disentanglement of diffusion frequency-dependence, tensor shape, and relaxation using multidimensional MRI

**DOI:** 10.1101/2023.10.10.561702

**Authors:** Jessica T.E. Johnson, M. Okan Irfanoglu, Eppu Manninen, Thomas J. Ross, Yihong Yang, Frederik B. Laun, Jan Martin, Daniel Topgaard, Dan Benjamini

## Abstract

Diffusion MRI with free gradient waveforms, combined with simultaneous relaxation encoding, referred to as multidimensional MRI (MD-MRI), offers microstructural specificity in complex biological tissue. This approach delivers intravoxel information about the microstructure, local chemical composition, and importantly, how these properties are coupled within heterogeneous tissue containing multiple microenvironments. Recent theoretical advances incorporated diffusion time dependency and integrated MD-MRI with concepts from oscillating gradients. This framework probes the diffusion frequency, *ω*, in addition to the diffusion tensor, **D**, and relaxation, *R*_1_, *R*_2_, correlations. A **D**(*ω*)-*R*_1_-*R*_2_ clinical imaging protocol was then introduced, with limited brain coverage and 3 mm^3^ voxel size, which hinder brain segmentation and future cohort studies. In this study, we introduce an efficient, sparse *in vivo* MD-MRI acquisition protocol providing whole brain coverage at 2 mm^3^ voxel size. We demonstrate its feasibility and robustness using a well-defined phantom and repeated scans of five healthy individuals. Additionally, we test different denoising strategies to address the sparse nature of this protocol, and show that efficient MD-MRI encoding design demands a nuanced denoising approach. The MD-MRI framework provides rich information that allows resolving the diffusion frequency dependence into intravoxel components based on their **D**(*ω*)-*R*_1_-*R*_2_ distribution, enabling the creation of microstructure-specific maps in the human brain. Our results encourage the broader adoption and use of this new imaging approach for characterizing healthy and pathological tissues.

## 1 INTRODUCTION

Several MRI approaches are inherently sensitive to microstructural features on the order of micrometers, probing cell morphology and organization, though, currently, they contend with having poor spatial resolution. Notably, diffusion MRI (dMRI) (Stejskal & Tanner 1965) is an important tool that encodes diffusive water displacement, which is sensitive to cell membranes and physical barriers (Beaulieu 2002; Leuze et al. 2017; Williamson et al. 2019), making it well-suited for indirectly assessing sizes and shapes of tissue components (Basser, Mattiello, & LeBihan 1994; Burcaw, Fieremans, & Novikov 2015; Komlosh et al. 2018; LeBihan 1990; Novikov, Fieremans, Jespersen, & Kiselev 2019; Pierpaoli, Jezzard, Basser, Barnett, & Chiro 1996). Relaxometry MRI is a complementary approach that encodes temporal magnetization decay (e.g., longitudinal and transverse relaxation rates, *R*_1_ and *R*_2_, respectively), which allows it to probe different water microenvironments and is especially sensitive to the local chemical composition, such as the presence and volume fraction of macromolecules (Bouhrara et al. 2020; Dvorak et al. 2021; Labadie et al. 2014; Mackay et al. 1994).

Emergent diffusion-relaxation multidimensional MRI (MD-MRI) acquisition protocols, combining the best elements of these approaches by encoding both diffusion and relaxation “dimensions” simultaneously (Hurlimann, Burcaw, & Song 2006; Pizzolato et al. 2020; Stanisz & Henkelman 1998), have become a focal point in the field. This imaging method introduces additional information, namely, the correlations between diffusivity and relaxation, invaluable to the study of tissue microstructure (Benjamini & Basser 2017; Lundell et al. 2019; C. M. W. Tax 2020; Veraart, Novikov, & Fieremans 2018), brain connectivity (Benjamini & Basser 2020; DeSantis, Barazany, Jones, & Assaf 2016; Slator et al. 2021), and pathology (Benjamini et al. 2021; Kim, Doyle, Wisnowski, Kim, & Haldar 2017; Martin et al. 2021). Using diffusion acquisition schemes with free gradient waveforms (Sjölund et al. 2015) allows one to explore both the frequency-dependent and tensorial aspects of the encoding spectrum **b**(*ω*) (Lasič et al. 2022; Lundell & Lasic 2020), enabling the investigation of frequency/time-dependent changes of diffusion-relaxation correlations measures using a single framework (Narvaez, Svenningsson, Yon, Sierra, & Topgaard 2022).

Recent major advances in clinical translation (de Almeida Martins et al. 2020; Reymbaut et al. 2021) led to an implementation of a **D**-*R*_1_-*R*_2_ imaging protocol on a clinical scanner comprised of 633 volumes in total, with varying diffusion weightings and directions, tensor ranks, echo times, and repetition times (Martin et al. 2021). However, despite its potential, this 25-minute protocol did not provide full brain coverage (5 axial slices) and had a relatively large 3 mm^3^ voxel size. In addition, a proof-of-concept **D**-*R*_1_-*R*_2_ imaging protocol comprised of 134 volumes and 30 slices with 3 mm^3^ voxel size was recently demonstrated (Yon et al. 2023). Nevertheless, whole brain coverage and higher spatial resolution are imperative for applications that requires identification and segmentation of brain regions, thus enabling robust cohort studies. Therefore, designing a more efficient and sparse acquisition protocol is desirable.

Another under-explored area in MD-MRI is mitigating imaging artifacts due to the simultaneous use of multiple echo times and diffusion weightings. These artifacts, if not addressed properly, can impact data analysis, reliability, and reproducibility. Echo planar imaging (EPI), commonly used for collecting diffusion MRI (dMRI) data, introduces its own set of artifacts (Pierpaoli 2010; C. M. Tax, Bastiani, Veraart, Garyfallidis, & Irfanoglu 2022), with the signal-to-noise ratio (SNR) being a particular concern (Jones 2010). Denoising methods are therefore crucial for mitigating this SNR limitation and enhancing dMRI data analysis. Among various denoising techniques (Coupe et al. 2008; Knoll, Bredies, Pock, & Stollberger 2011; Tian et al. 2022), Marchenko-Pastur principal component analysis (MPPCA) has gained popularity for its ability to reduce noise without sacrificing anatomical detail or introducing blurriness, making it an effective approach for boosting SNR (Veraart, Novikov, et al. 2016). A recent advancement in denoising techniques, tensor-MPPCA (tMPPCA), extends the applicability of MPPCA to high-dimensional data (Olesen, Ianus, Østergaard, Shemesh, & Jespersen 2023). However, tMPPCA requires the multi-dimensional data to be densely acquired using a grid-like encoding, where, for example, there is an equal number of b-values and gradient directions for each echo time. Therefore, tMPPCA is not suitable for sparsely encoded multidimensional data that is used in the present study.

In this study, we first present an efficient and sparse *in vivo* frequency dependent MD-MRI acquisition protocol that provides whole brain coverage at 2 mm^3^ resolution. We demonstrate the feasibility and robustness of this pipeline using a well-defined phantom (Laun, Huff, & Stieltjes 2009) and repeated scans of five healthy participants. Second, to explore noise effects in the data, we assert that while MPPCA exploits the inherent redundancy in dMRI data (Veraart, Fieremans, & Novikov 2016), the sparse nature of the efficient MD-MRI encoding design cannot be considered highly redundant, and may require a more nuanced denoising strategy. Thus, we aimed to evaluate various denoising approaches and their impact on the reliability of outputs of the current MD-MRI framework. Analyzing the agreement between voxelwise **D**(*ω*)-*R*_1_-*R*_2_ distributions across two scanning sessions in the human brain, we compactly quantify similarities across high-dimensional spectra between corresponding voxels using the earth mover’s distance (EMD) (Rubner, Tomasi, & Guibas 1998).

## 2 MATERIALS AND METHODS

### 2.1 Phantom

We used an anisotropic phantom (HQ imaging, Lörrach, Germany) to assess the robustness of our MD-MRI acquisition and processing pipeline, and the effects of different denoising strategies. The basic properties of this phantom have previously been described in detail (Laun, Huff, & Stieltjes 2009). Briefly, the phantom consists of parallel polyamide fibers of 15 *μ*m diameter wound on a circular polyoxymethylene spindle. An aqueous solution was used as embedded fluid between the fibers, resulting in restricted diffusion perpendicular to the fibers. The spindle with the fibers was immersed in an aqueous polyvinylpyrrolidone (PVP) solution (Pierpaoli, Sarlls, Nevo, Basser, & Horkay 2009; Wagner et al. 2017). This design creates two regions of interest (ROIs): fluid and tightly packed fibers, with isotropic and anisotropic diffusion characteristics, respectively.

### 2.2 Participants

Five healthy participants (ages 41.2 ± 8.3, 3 women) were each scanned twice, a few weeks apart (i.e., total of 10 scans). Participants were systematically drawn from ongoing healthy cohorts of the National Institute on Drug Abuse (NIDA). Experimental procedures were performed in compliance with our local Institutional Review Board, and participants provided written informed consent. Prior to each scan, NIDA clinical and nursing units conducted Covid-19 testing, urine drug tests, a physical exam, and a questionnaire on pre-existing conditions and daily habits. Exclusion criteria included major medical illness or current medication use, a history of neurological or psychiatric disorders or substance abuse.

### 2.3 Data acquisition

Phantom and human data were acquired using a 3T scanner (MAGNETOM Prisma, Siemens Healthcare AG, Erlangen, Germany) with a 32 channel head coil. Data were acquired with 2 mm isotropic voxel size using a single-shot spin-echo EPI sequence (Wetscherek, Stieltjes, & Laun 2015) modified for tensor-valued diffusion encoding with free gradient waveforms (Martin et al. 2021). The acquisition parameters were set as follows: FOV = 228 × 228 × 110 mm^3^, voxel size = 2 × 2 × 2 mm^3^, acquisition bandwidth = 1512 Hz/Px, in-plane acceleration factor 2 using GRAPPA reconstruction with 24 reference lines, effective echo spacing of 0.8 ms, phase-partial Fourier factor of 0.75, and axial slice orientation.

The acquisition protocol was designed following previously described heuristic guidelines (Martin et al. 2021). A detailed summary of the acquisition protocol is displayed in Fig. 1 . In short, in addition to a *b* = 0 ms/*μ*m^2^ volume, numerically optimized (Sjölund et al. 2015) linear, planar, and spherical b-tensors were employed with b-values ranging between 0.1 and 3 ms/*μ*m^2^. Here we augment the data acquisition scheme with exploration of the *ω*-dimension of **b**(*ω*) in the range of 6.6 - 21 Hz centroid frequencies, *ω*_*cent*_/2*π*, to allow the decoupling of frequency-dependent diffusion components. It should be noted that the highest frequencies reached for b-values of 0.5, 1.5, and 3 ms/*μ*m^2^ were 21, 15, and 11 Hz, respectively. The datasets were acquired with a single phase encoding direction (anterior to posterior, AP), and an additional *b* = 0 ms/*μ*m^2^ volume with reversed phase encoding direction (PA). Sensitivity to *R*_1_ and *R*_2_ was achieved by acquiring data with different combinations of repetition times, TR=(0.62, 1.75, 3.5, 5, 7, 7.6) s and echo times, TE=(40, 63, 83, 150) ms. The number of concatenations and preparation scans was increased to allow values of TR below 5 s. To achieve an efficient and sparse acquisition, encoding parameters that maximize signal differences between brain tissue types, while maintaining sufficient SNR, were empirically selected. A total of 139 individual measurements were recorded over 40 min.

**FIGURE 1.**
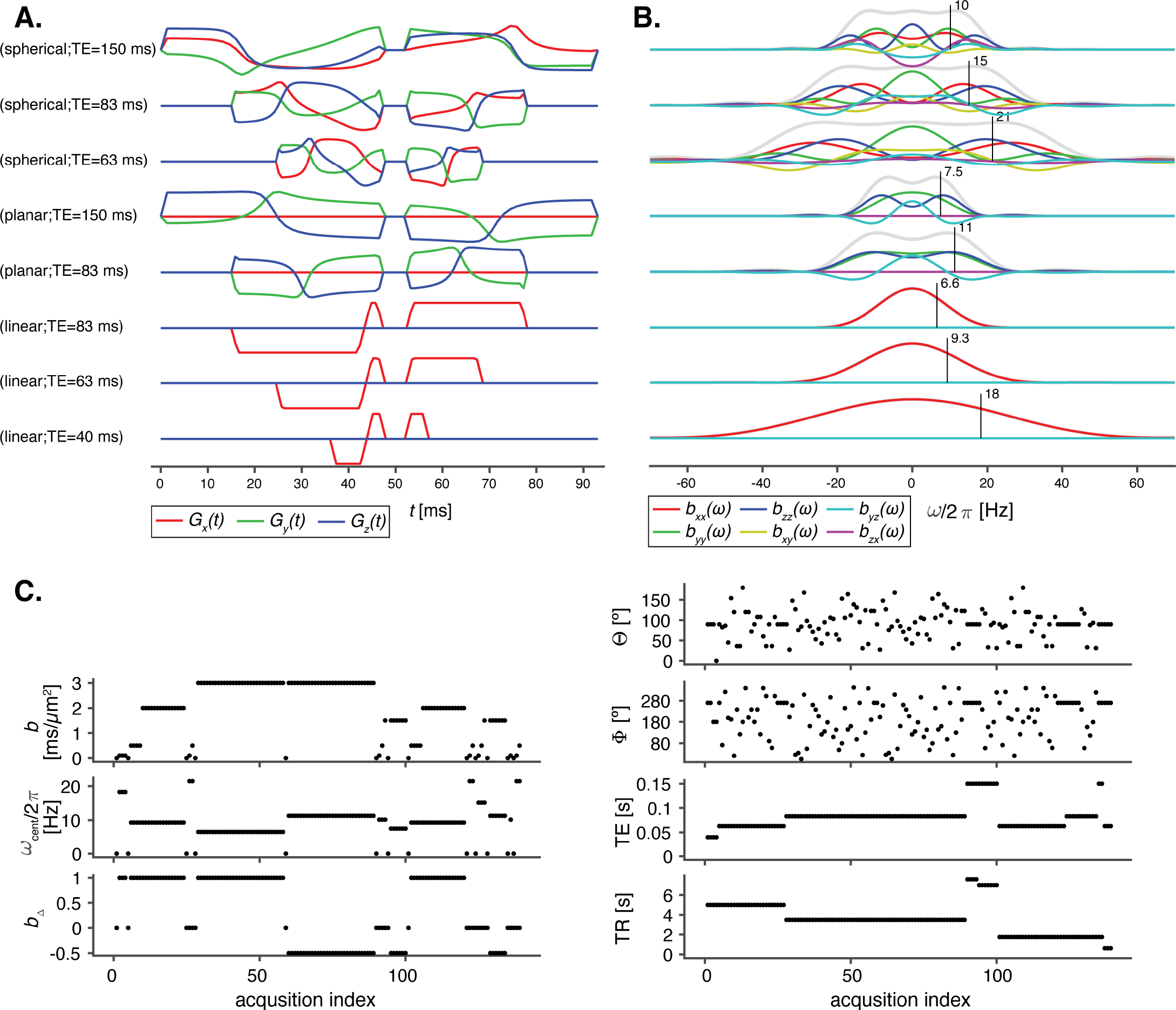
Key experimental details. (A) Time-dependent effective gradients *G*(*t*) and (B) corresponding tensor-valued encoding spectra **b**(*ω*) for linear, planar, and spherical encoding at different echo times and centroid frequencies, denoted by black vertical lines. (C) Acquisition protocol with repetition time TR, echo time TE, as well as **b**-tensor magnitude *b*, normalized anisotropy *b*_Δ_ (planar: –0.5, spherical: 0, linear: 1), orientation (Θ, Φ), and centroid frequency *ω*_*cent*_/2*π*, versus image acquisition index.

### 2.4 Preprocessing and denoising strategies

Two denoising strategies were evaluated, in which MPPCA was applied on (1) the combined MD-MRI data and on (2) MD-MRI data grouped according to TE. These were compared with skipping the denoising step altogether and with a reference pipeline without any processing steps. The four strategies are schematically shown in Fig. 2 .

**FIGURE 2.**
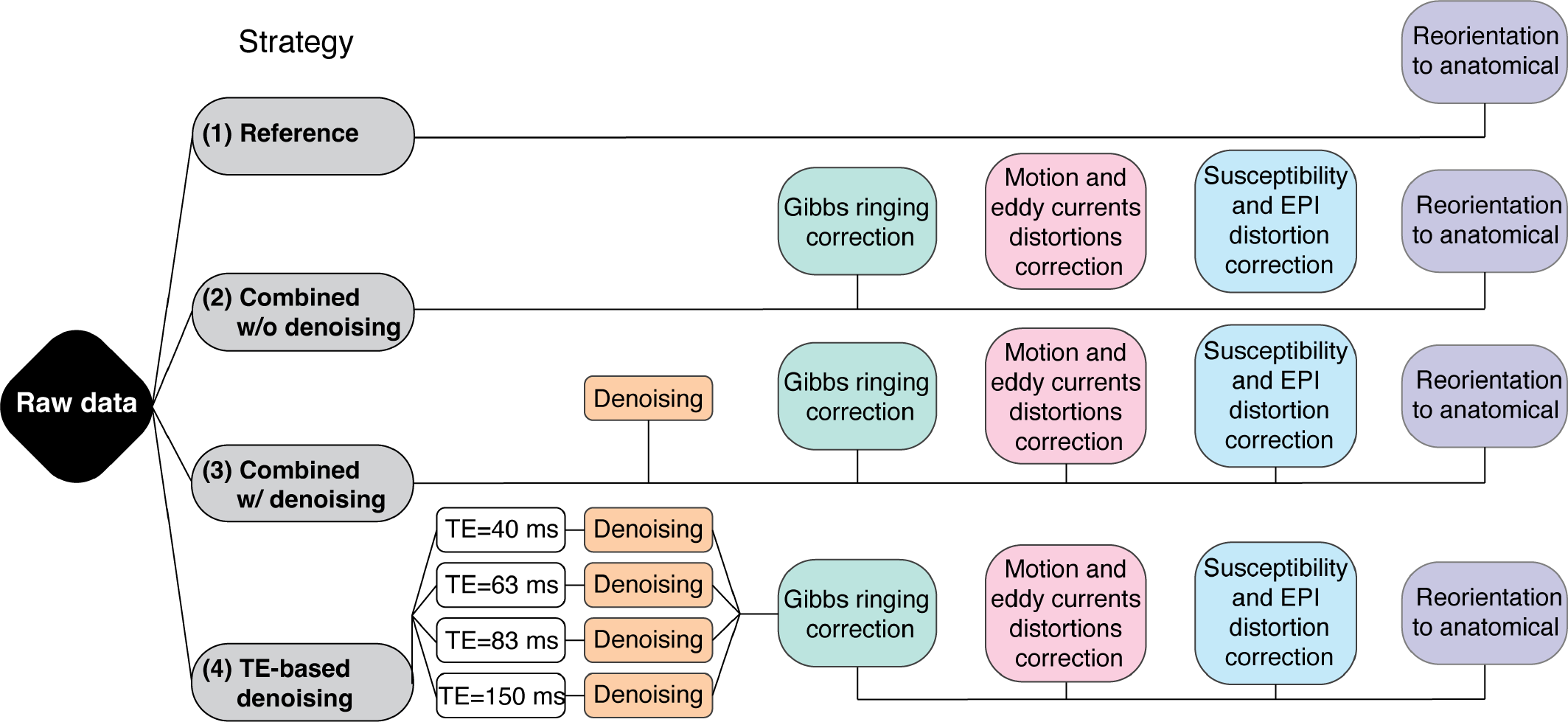
Schematic description of the evaluated denoising strategies. The Reference pipeline did not include any processing steps besides reorientation to the anatomical image space. In Strategies 2 and 3, all volumes, regardless of TE/TR and b-tensor encoding design, were initially combined into a single dataset. The MPPCA denoising step was skipped in Strategy 2, and turned on in Strategy 3. For Strategy 4, all datasets were grouped according to echo time prior to denoising. The grouped data were then combined again after denoising for the remainder of the pipeline.

The preprocessing modules used in this work are part of the TORTOISE dMRI processing package (M. Irfanoglu, Nayak, Taylor, & Pierpaoli 2023; Pierpaoli et al. 2010). For the full pipeline, the dMRI data initially underwent denoising with the MPPCA technique (Veraart, Novikov, et al. 2016), which was followed by Gibbs ringing correction (Kellner, Dhital, Kiselev, & Reisert 2016) for partial k-space acquisitions (H. Lee, Novikov, & Fieremans 2021). Motion and eddy currents distortions were subsequently corrected with TORTOISE’s DIFF-PREP module (Rohde, Barnett, Basser, Marenco, & Pierpaoli 2004) with a physically-based parsimonious quadratic transformation model and a normalized mutual information metric. For susceptibility distortion correction, a T1W image was initially converted to a T2W image with b=0 s/mm^2^ contrast (Schilling et al. 2019), which was fed into the DRBUDDI (M. O. Irfanoglu et al. 2015) method for AP PA distortion correction. The final preprocessed data was output with a single interpolation in the space of an anatomical image at native in-plane voxel size. We note that the data virtually did not contain any slice-to-volume motion or motion-induced signal dropouts, therefore, these options were disabled in the processing.

We applied the original recommendation of choosing the patch size for MPPCA denoising so that the resulting matrices are approximately square (Veraart, Novikov, et al. 2016). When denoising the combined dataset, which consists of 139 volumes, the patch size was [5×5×5] voxels. When denoising data grouped according to TE (echo times of 40, 63, 83, 150 ms), which consists of 4, 49, 73, 13 volumes, respectively, the patch sizes were [2×2×2], [4×4×4], [4×4×4], and [2×2×2] voxels, respectively.

### 2.5 Multidimensional data processing

The preprocessed data were processed in Matlab R2019b (MathWorks, Natick, MA) using the Monte Carlo inversion algorithm (Narvaez et al. 2022) as implemented in the multidimensional diffusion MRI toolbox (Nilsson et al. 2018). Briefly, the **b**(*ω*) − TE − TR encoded signal *S* is modeled as a sum of contributions; the *i*th component is characterized by its signal weight, or fraction *f*_*i*_, tensor-valued diffusion spectra **D**_*i*_(*ω*), and longitudinal and transverse relaxation rates *R*_1,*i*_ and *R*_2,*i*_ according to (Narvaez et al. 2022)

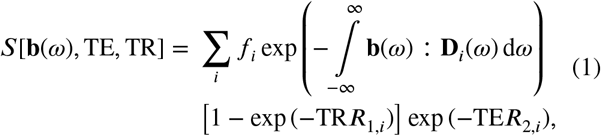

where the colon denotes a generalized scalar product.

As previously described in detail (Narvaez et al. 2021), inversion of Eq. 1 is rendered tractable by approximating **D**_*i*_(*ω*) as axisymmetric Lorentzians parameterized by the zero-frequency axial and radial diffusivities [*D*_||,*i*_, *D*_⊥,*i*_], azimuthal and polar angles [Θ_*i*_, Φ_*i*_], high frequency isotropic diffusivity *D*_0,*i*_, axial and radial transition frequencies, [Γ_||,*i*_, Γ_⊥,*i*_], along with longitudinal and transversal relaxation rates [*R*_1,*i*_, *R*_2,*i*_]. In this study, these parameters were sampled in the ranges 0.05 ≤ *D*_||/⊥/0_ ≤ 5 *μ*m^2^/ms, 0 ≤ Θ ≤ *π*, 0 ≤ Φ ≤ 2*π*, 0.2 ≤ *R*_1_ ≤ 2 s^−1^, 1 ≤ *R*_2_ ≤ 30 s^−1^, and 0.01 ≤ Γ_||/⊥_ ≤ 10000 s^−1^. The Monte Carlo inversion algorithm finds an ensemble of solutions within the above sampling range, and estimates the corresponding weights *f* via nonnegative least-squares, iterating this process while applying quasi-genetic filtering and bootstrapping with replacement to account for the inherent ill-conditioned nature of the Laplace inversion (Benjamini 2020; de Almeida Martins & Topgaard 2018). Following the terminology in (de Almeida Martins & Topgaard 2018), the Monte Carlo inversion was performed with *N*_*in*_ = 200 input components, *N*_*p*_ = 20 proliferation steps, *N*_*m*_ = 20 mutation steps, *N*_*out*_ = 10 output components, and *N*_*b*_ = 100 rounds of bootstrapping.

Voxelwise **D**(*ω*)-*R*_1_-*R*_2_ distributions in the primary analysis space [*D*_||_, *D*_⊥_, Θ, Φ, *D*_0_, Γ_||_, Γ_⊥_, *R*_1_, *R*_2_] were evaluated at selected values of *ω* within the narrow 6.6 - 21 Hz window actually probed by the gradient waveforms, giving a set of *ω*-dependent distributions in the [*D*_||_ (*ω*), *D*_⊥_ (*ω*), Θ, Φ, *R*_1_, *R*_2_] space. For each value of *ω*, the results are visualized as *ω*-independent distributions by projecting *D*_||_ (*ω*) and *D*_⊥_(*ω*) to the dimensions of isotropic diffusivity *D*_*iso*_(*ω*) and squared normalized anisotropy 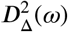 (Conturo, McKinstry, Akbudak, & Robinson 1996), according to

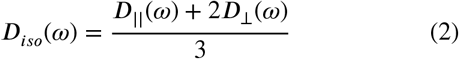

and

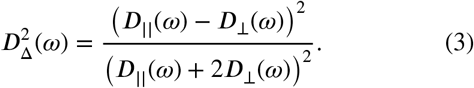

The means, variances, and covariances over relevant dimensions and sub-divisions (“bins”) of the distribution space are then computed. There are typically three bins in the *in vivo* human brain, roughly corresponding to white matter (WM), gray matter (GM), and CSF (Martin et al. 2021). These *in vivo* bins, which will be respectively referred to as bin 1, 2, and 3, represent partial integration regions in the 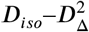 distribution space, i.e., bin 1: *D*_*iso*_ < 2.5*μ*m^2^/ms and 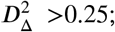 bin 2: *D*_*iso*_ < 2.5*μ*m^2^/ms and 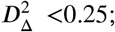 and bin 3: *D*_*iso*_ > 2.5*μ*m^2^/ms and the full range of 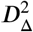. The normalized weights of these bins were mapped and are labeled as *f*_*bin*1_, *f*_*bin*2_, and *f*_*bin*3_. The phantom we used has two distinct regions, isotropic and anisotropic diffusion, and therefore only two bins were used, with the following partial integration regions in the 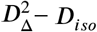 distribution space: *D*_*iso*_ < 1.25*μ*m^2^/ms and 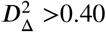 for bin 1 (anisotropic), and *D*_*iso*_ > 1.15*μ*m^2^/ms and 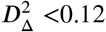 for bin 2 (isotropic). The normalized weights of these bins were mapped and are labeled as *f*_*aniso*_ and *f*_*iso*_. Following conventions often used to display results from oscillating gradient encoding (Aggarwal, Jones, Calabresi, Mori, & Zhang 2012) the effects of restricted diffusion were quantified by a finite difference approximation of the rate of change of the diffusivity metrics with frequency within the investigated window, which in our case was 21 Hz and 6.6 Hz.

### 2.6 Error estimates from phantom measurements

The ROIs from the isotropic and anisotropic portions of the phantom were used to assess the effect of different denoising strategies on the error estimate from the MD-MRI pipeline. We note that the signal intensity has been estimated from the fitted models; one hundred bootstrap solutions have been estimated, the expected signal intensity for each measurement has been estimated from the models, and the median over those signal estimates has been taken. Let *y*_*m,v*_ be the measured normalized signal intensity of measurement (i.e., image volume) *m* at voxel *v*. Let ŷ_*m,v*_ be the estimated signal intensity.

We then compute the root-mean-square error (RMSE) for each measurement and for each voxel. To compute the RMSE for each measurement, RMSE_*m*_, we square the difference between the measured and fitted voxel-wise normalized signal for each measurement and voxel, average those squared differences over the voxels and take the square root:

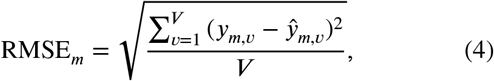

where *V* is the number of voxels.

To obtain the RMSE for each voxel, RMSE_v_ , (i.e., error for the whole fit instead of an error for each measurement), we compute

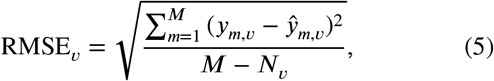

where (*M* − *N*_*v*_) is the error degrees of freedom, *M* is the number of measurements, and *N*_*v*_ is the number of estimated parameters at voxel *v*. The median value of *N*_*v*_ over the bootstraps is used because *N*_*v*_ can be different for each bootstrap.

### 2.7 Variability analysis

The effect of different denoising strategies on the variability of the MD-MRI estimates was quantified by computing the distance between pairs of corresponding (i.e., voxelwise) **D**(*ω*)-*R*_1_-*R*_2_ distributions from the first and second scans within each participant. These computations were performed in the midway space between the scan and rescan volumes for each participant space using a rigid transformation to ensure that the image registration process exerted an equivalent influence on the scan and rescan data (Reuter, Schmansky, Rosas, & Fischl 2012; Veraart, Raven, Edwards, Weiskopf, & Jones 2021). This midway transformation was applied to the discrete **D**(*ω*)-*R*_1_ -*R*_2_ coordinates [*D*_||_(*ω*), *D*_⊥_(*ω*), Θ, Φ, *R*_1_, *R*_2_] and their respective weights in each imaging session’s native space, bringing each map to the midway point between the scan and rescan sessions.

We propose here the Earth Mover’s Distance (EMD) (Rubner et al. 1998) as a novel distance measure between **D**(*ω*)-*R*_1_-*R*_2_ distributions, once both scan sessions have been transformed to the midway space. In the current context, the EMD between voxelwise distributions obtained from two scans from the same subject will produce a complete description of the variability using different denoising strategies. The EMD computes the distance between two distributions of any dimension, expressed as discrete coordinates and weights (as in the current implementation), or as density functions (continuous representations, e.g., (Benjamini et al. 2020)), which are represented by signatures. The signatures are sets of weighted features that capture the distributions. In our case, the features were the discrete **D**(*ω*)-*R*_1_ -*R*_2_ coordinates [*D*_||_ (*ω*), *D*_⊥_(*ω*), Θ, Φ, *R*_1_, *R*_2_].

As described in 2.5, each voxel contains *N*_*b*_ = 100 bootstrap estimates of **D**(*ω*)-*R*_1_-*R*_2_. Therefore, all discrete coordinates within each bootstrap iteration, [*D*_||_(*ω*), *D*_⊥_(*ω*), Θ, Φ, *R*_1_, *R*_2_]_*n*_, and their respective weights, *f*_*n*_, were joined together, followed by a final normalization step of the weights. The EMD is defined as the minimum amount of “work” needed to change one signature into the other. The work is based on predefined ground distance, i.e., the distance between two features in corresponding voxels, which was chosen here to be Euclidean distance. Because of their different magnitudes and units, the features, [*D*_||_(*ω*), *D*_⊥_(*ω*), Θ, Φ, *R*_1_, *R*_2_], were normalized according to their respective maximal values, detailed in 2.5. Further, the angles [Θ, Φ] were wrapped to [*π*, 2*π*], respectively, to take the periodicity and inversion symmetry of the orientation space into account.

### 2.8 Statistical analysis

Voxelwise RMSE (RMSE_*v*_) from the isotropic and anisotropic ROIs within the phantom were grouped according to denoising strategy and compared using one-way ANOVA. In a similar manner, voxelwise EMD from the five subjects were first joined together, grouped according to denoising strategy, and compared using one-way ANOVA. In both phantom and *in vivo* cases, a multiple comparison test using the Bonferroni correction was performed to determine the effect the different denoising strategies have on the MD-MRI estimates.

## 3 RESULTS

### 3.1 Diffusion phantom

We first assessed various denoising approaches based on the RMSE of the MD-MRI model fit, considering both isotropic and anisotropic ROIs within the phantom. It is important to note that RMSE serves as a measure of the model’s fidelity in capturing the measurements, although it may not independently validate the model. However, when a straightforward physical ground truth is available that is expected to be well-described by the MD-MRI model, as in the case of the phantom, RMSE serves as a robust indicator of the SNR. Figure 3 A shows the RMSE of each measurement averaged over all voxels within each ROI, for each denoising strategy. Especially notable in the isotropic ROI, some volumes have higher fit errors than others, which may indicate more severe image artefacts. However, the RMSE_*m*_ remains below 1% of the *S*_0_ signal intensity for all except two volumes in the isotropic ROI, and below 3% in the anisotropic ROI.

**FIGURE 3.**
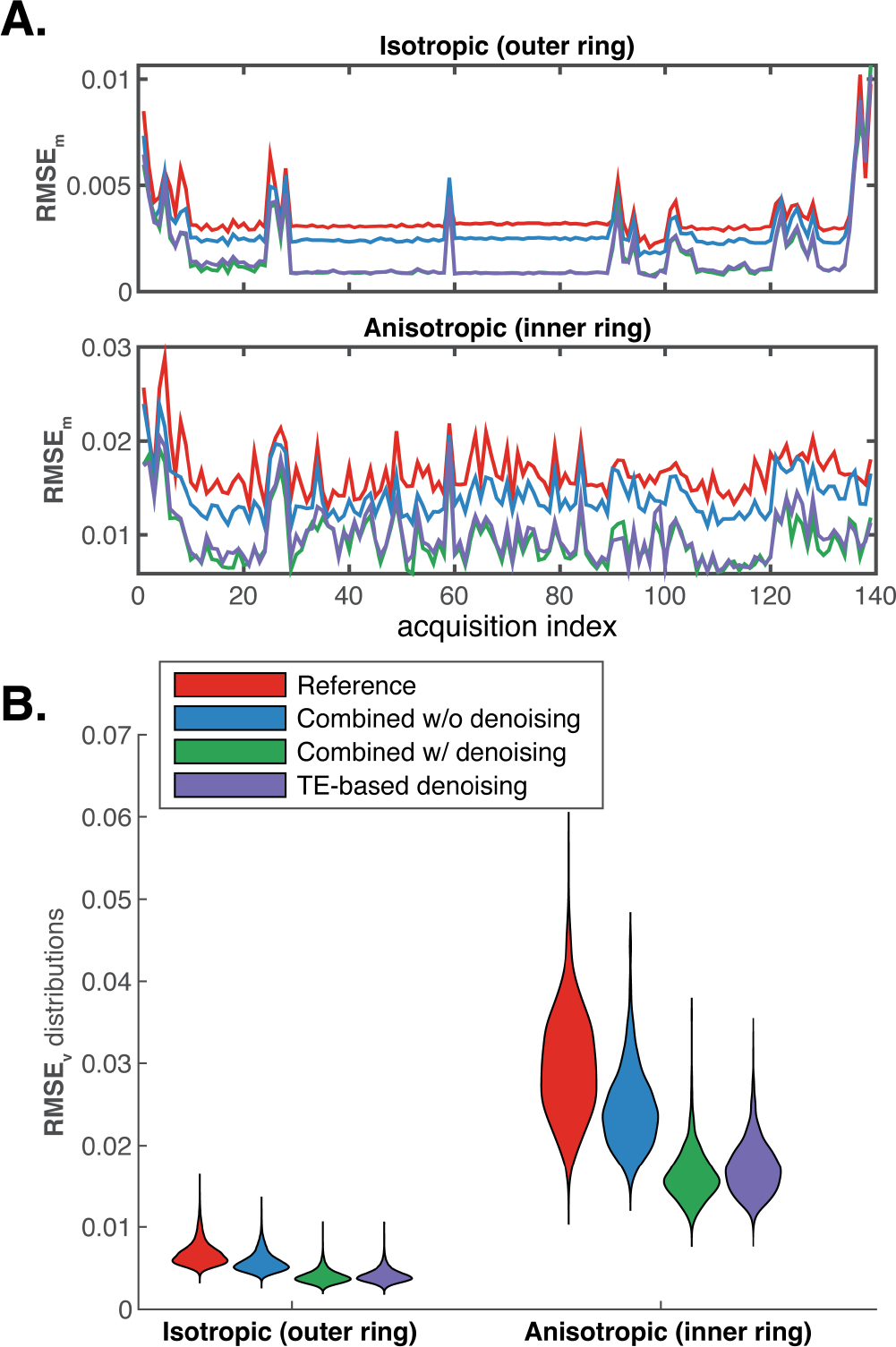
Effect of denoising strategy on the model fit root-mean-square error (RMSE). (A) Normalized RMSE for each measurement (image volume) averaged over the isotropic (top) and anisotropic (bottom) ROIs, color coded according to denoising strategies. (B) Distribution of normalized RMSE over voxels within each ROI, color coded according to denoising strategies.

Voxelwise RMSE distributions for each ROI and denoising strategy are presented in Fig. 3 B. One-way ANOVA in the isotropic ROI showed that the source of most mean squares variability was between groups (*p* < 0.00001), with RMSE_*v*_ means of 6.75e-3, 5.72e-3, 3.95e-3, and 4.06e-3, for the Reference, combined data without and with denoising, and TE-based denoising, respectively. Pairwise comparisons using a multiple comparison test with the Bonferroni method identified that all the denoising strategies have significantly different RMSE_*v*_ means. Similar analysis was done with the anisotropic ROI voxels, with RMSE_*v*_ means of 2.98e-2, 2.45e-2, 1.63e-2, and 1.72e-2, for the Reference, combined data without and with denoising, and TE-based denoising, respectively. Here too, most mean squares variability was due to differences among the group means (*p* < 0.00001), and Bonferroni-corrected pairwise comparisons showed that all the denoising strategies have significantly different RMSE_*v*_ means. These results demonstrate an advantage towards combining the sparse MD-MRI dataset first, and then applying MPPCA denoising on all the volumes at once.

Next, we illustrate the stability of our efficient MD-MRI acquisition protocol using a well-defined anisotropic phantom. Figure 4 A shows the way in which sub-voxel microstructural information is obtained by partitioning the 2D 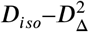 plane into two regions (or bins), based on isotropic diffusion length scale and diffusion anisotropy. In the case of the phantom, in which only two well-defined diffusion populations exist, applying the partial integration in Fig. 4 A in each voxel results in signal fraction map (*f*_*aniso*_,*f*_*iso*_) coded into red and blue colors, respectively, and shown in Fig. 4 B. Single-voxel attenuation profiles (colored circles) and their fits (black dots) from both ROIs are shown in Fig. 4 C.

**FIGURE 4.**
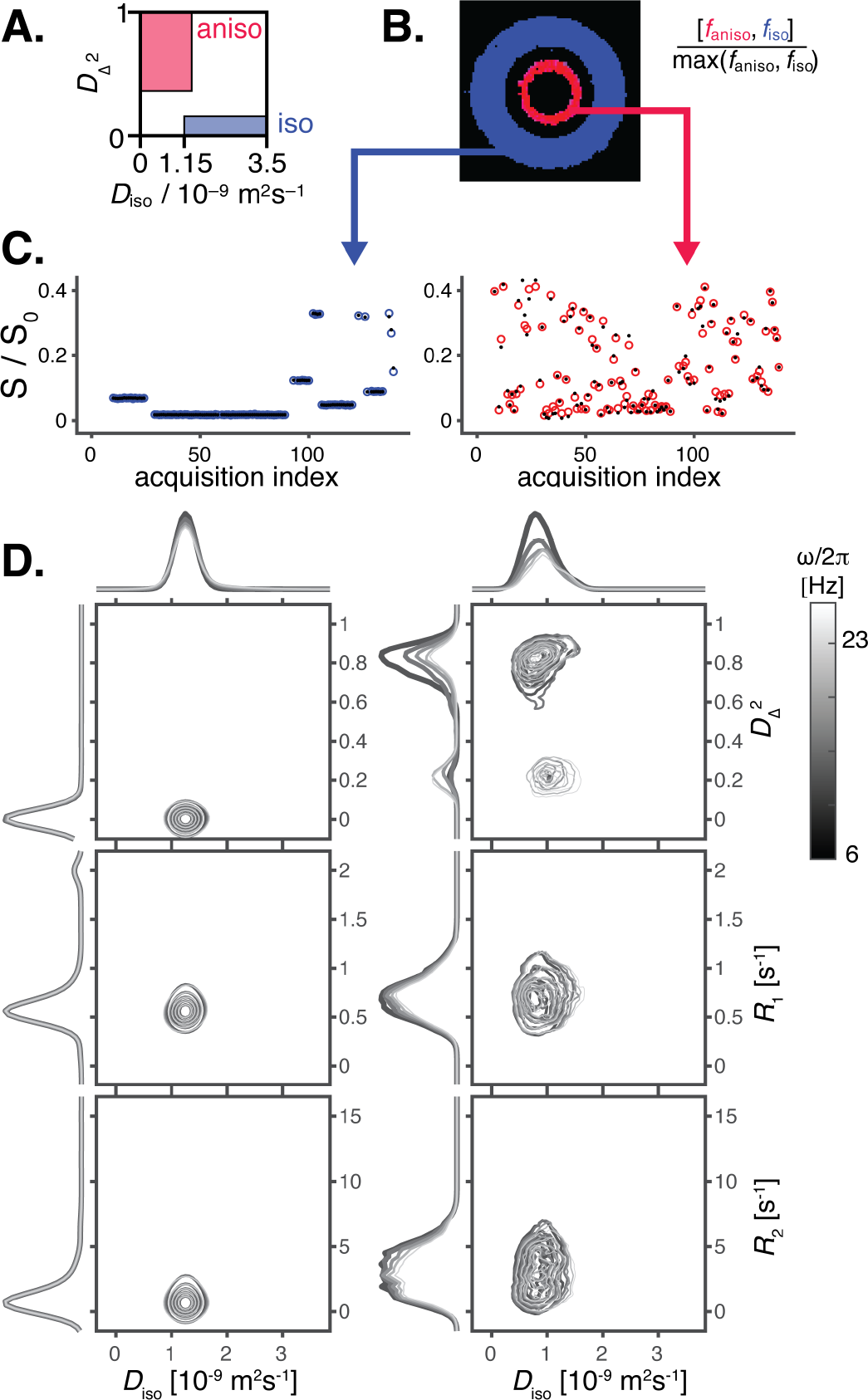
Representative phantom voxels from the isotropic (blue) and anisotropic (red) ROIs. (A) Bin segmentation between the two components representing partial integration regions in the 2D 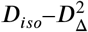 plane. (B) The resulting isotropic and anisotropic signal fraction maps, color-coded (blue=isotropic, red=anisotropic). (C) Single-voxel attenuation profiles (colored circles) and their fits (black dots). (D) **D**(*ω*)-*R*_1_-*R*_2_ distributions for each voxel projected onto the 2D 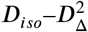, *D*_*iso*_–*R*_1_, and *D*_*iso*_–*R*_2_ planes for five frequencies in the range of *ω*/2*π*= 6.6-21 Hz as indicated with the linear gray scale of the contour lines.

The per-voxel **D**(*ω*)-*R*_1_-*R*_2_ distributions from both the isotropic and anisotropic ROIs are visualized in Fig. 4 D as projections onto the 2D 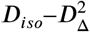, *D*_*iso*_–*R*_1_, and *D*_*iso*_–*R*_2_ planes for frequencies *ω*/2*π* between 6.6 and 21 Hz (represented by the grayscale intensity of the contour plots). The microstructural differences between the ROIs are most visible in the 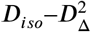 projection from a representative single voxel, in which water within the isotropic ROI appears to experience completely isotropic diffusion (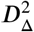 close to 0), and *D*_*iso*_ of about 1.25 *μ*m^2^/ms, without any notable diffusion frequency dependency. Mostly single peaks in the *D*_*iso*_ –*R*_1_ and *D*_*iso*_–*R*_2_ planes point to low relaxation rates, as expected from aqueous PVP solution (Pierpaoli et al. 2009). The 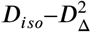 projection from a single voxel within the anisotropic diffusion ROI shows pronounced diffusion frequency/time dependence behavior. A single water population with low *D*_*iso*_ of about 0.78 *μ*m^2^/ms and high 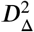 of about 0.82 is observed at low frequency. A microenvironment with the same *D*_iso_ but with lower apparent anisotropy (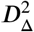 of about 0.20) is observed as the frequency increases. Although 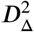 values of about 0.2 have been previously observed in the brain, corresponding to a nearly symmetric “butterfly” spread of components centered about the *D*_Δ_ = 0 line (de Almeida Martins et al. 2020; Yon et al. 2020), this inversion related artifact is not present in our data, as can be seen by the strictly positive *D*_Δ_ in Supplementary Fig. 1. Instead, the low anisotropy peak provides direct evidence of the coupling of diffusion length and time scales. This diffusion time/frequency dependency behavior in diffusion phantoms is expected when the asymptotic diffusion time is not reached, and the characteristic length scale, 7.6 *μ*m in our case, cannot be fully sampled. While at 6.6 Hz the diffusion length scale of about 11 *μ*m should fully sample the microstructure, at 21 Hz and a diffusion length scale of about 6 *μ*m, a proportion of the diffusing water is not fully restricted by the boundaries.

Figure 5 displays a compilation of parametric maps representing global and bin-specific statistical characteristics derived from the voxelwise **D**(*ω*)-*R*_1_-*R*_2_ distributions, preprocessed using all volumes combined for the denoising step. The axial slice of the phantom features two concentric rings: the inner ring, composed of 15 *μ*m diameter fibers in an aqueous solution, exhibits highly anisotropic microstructure, while the outer ring contains aqueous PVP solution, known for its isotropic properties.

**FIGURE 5.**
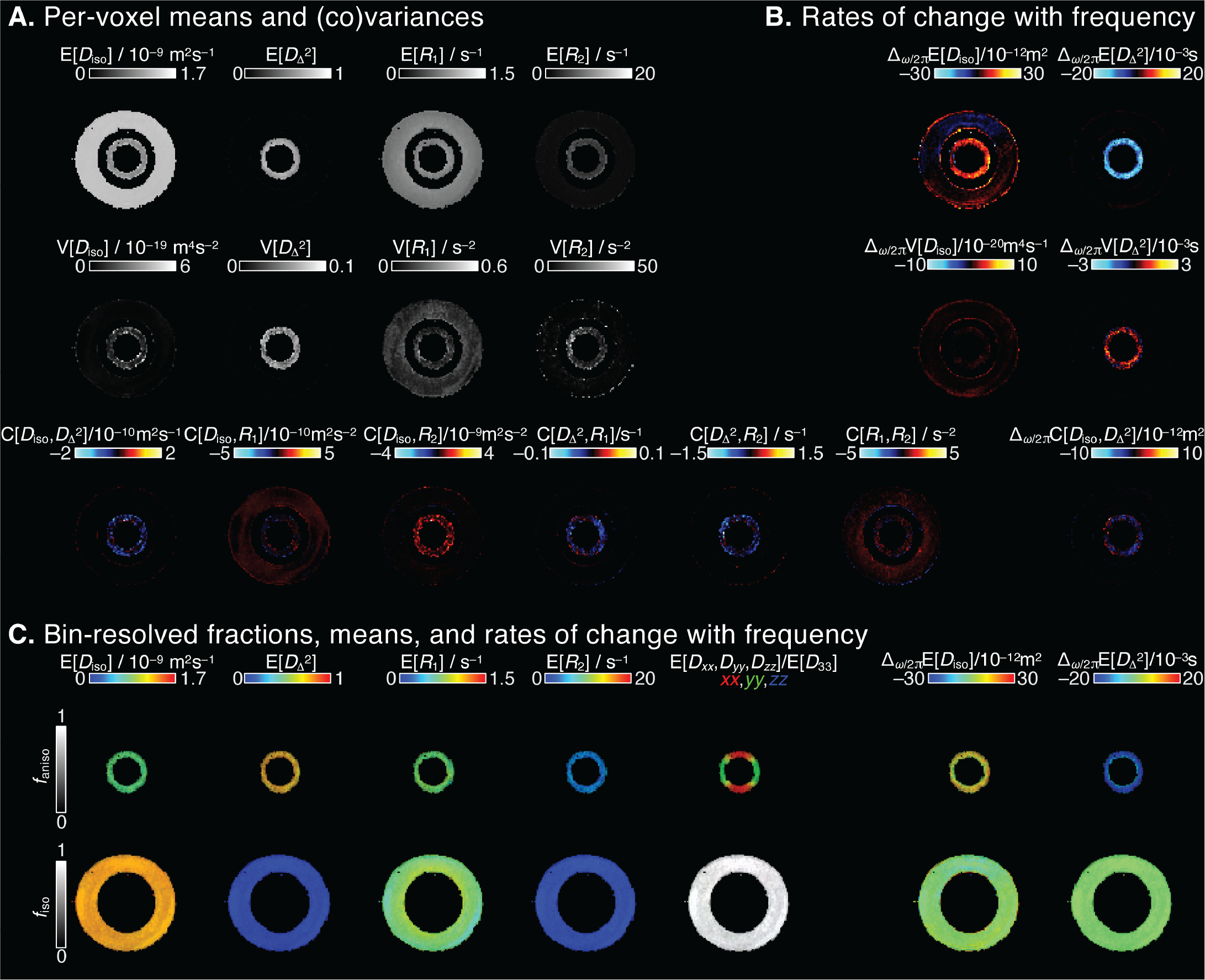
Parameter maps of the diffusion phantom derived from voxelwise **D**(*ω*)-*R*_1_-*R*_2_ distributions. (A) Voxelwise means E[*x*], variances V[*x*], and covariances C[*x, y*] at a selected encoding frequency *ω*/2*π* = 6.6 Hz. (B) Parameter maps of the rate of change with frequency, Δ_*ω*/2*π*_E[x]. (C) Bin-resolved maps of E[*x*] and Δ_*ω*/2*π*_E[x] according to Fig. 4 A. The brightness and color scales represent, respectively, the signal fractions and the values of each parameter.

In Fig. 5 A, the maps illustrate per-voxel statistics, including means E[*x*], variances V[*x*], and covariances C[*x, y*] for *D*_*iso*_, 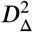, *R*_1_, and *R*_2_, assessed at *ω*/2*π* = 6.6 Hz. The means E [*D*_*iso*_], E[*R*_1_], and E[*R*_2_] correspond to conventional mean diffusivity (Basser et al. 1994), quantitative *R*_1_, and *R*_2_ (Weiskopf, Edwards, Helms, Mohammadi, & Kirilina 2021), respectively. The 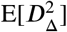 map is akin to metrics used to quantify microscopic diffusion anisotropy (Lasič, Szczepankiewicz, Eriksson, Nilsson, & Topgaard 2014; Lawrenz, Koch, & Finsterbusch 2010; Shemesh et al. 2015). Averaged across the ROIs, the E[*D*_*iso*_] values were 0.96±0.07 *μ*m^2^/ms and 1.29±0.01 *μ*m^2^/ms, for the anisotropic and isotropic ROIs, respectively, in agreement with previous measurements (Laun, Schad, Klein, & Stieltjes 2009). As expected, the microstructural difference between the ROIs is demonstrated from the averaged anisotropic and isotropic 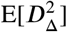 values of 0.63±0.05 and 0.004±0.002, respectively. The increased intravoxel variances in the anisotropic ROI compared to the isotropic region suggest potential heterogeneity and imperfect fiber packing.

Examining the relaxation properties of the phantom, we measured faster transverse relaxation rates in the anisotropic ROI (E[*R*_2_]=4.76±0.77 s^−1^ and 1.15±0.14 s^−1^, for the anisotropic and isotropic ROIs, respectively). This observation aligns with expectations of increased relaxation in the presence of polymer fibers due to surface relaxation effects (Laun, Huff, & Stieltjes 2009). In addition, we observed comparable longitudinal relaxation rates of E[*R*_1_]=0.76±0.05 s^−1^ and 0.74±0.06 s^−1^ in the anisotropic and isotropic ROIs, respectively.

The frequency-dependence results, presented in Fig. 5 B, show the rate of change with frequency within the investigated range of 6.6–21 Hz for the per-voxel means, variances, and covariance of *D*_*iso*_ and 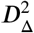. In the Δ_*ω*/2*π*_E[*D*_*iso*_] map, positive values indicate diffusion time dependency behavior suggestive of restriction (Aggarwal et al. 2012). Conversely, decreased anisotropy with higher frequency results in negative values in the 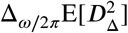 map. In both cases, the phantom regions demonstrated the expected results: positive and negative values of Δ_*ω*/2*π*_E[*D*_*iso*_] and 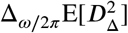 in the anisotropic ROI, respectively, and negligible frequency dependency in the isotropic ROI.

We can extract subvoxel information from the 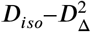 plane bins depicted in Fig. 4 A. These bin-resolved maps, displayed in Fig. 5 C, illustrate the means of the tensorial, relaxation, and frequency properties. Each map employs two orthogonal scales: the brightness intensity reflects the relative signal fraction, while the color scale represents the specific property value. In addition to these metrics, we computed the voxelwise principal orientation of the tensors and visualized it as directionally-encoded color (DEC) maps for each bin, utilizing the tensor orientation data (i.e., [Θ, Φ]). The DEC map pattern in Fig. 5 C aligns with findings from previous studies (Laun, Huff, & Stieltjes 2009; Laun, Schad, et al. 2009).

### 3.2 *In vivo* MD-MRI

We first assessed the impact of different denoising strategies on the MD-MRI pipeline and estimates. Voxelwise agreement between the first and second scan sessions was quantified using the EMD, as detailed in 2.7. We calculated EMD histograms for the entire brain averaged across the five subjects for each denoising method, as depicted in Fig. 6 A. A closer examination of the histogram highlights that denoising the combined MD-MRI data yields the lowest overall EMD values across the entire brain for the five subjects.

**FIGURE 6.**
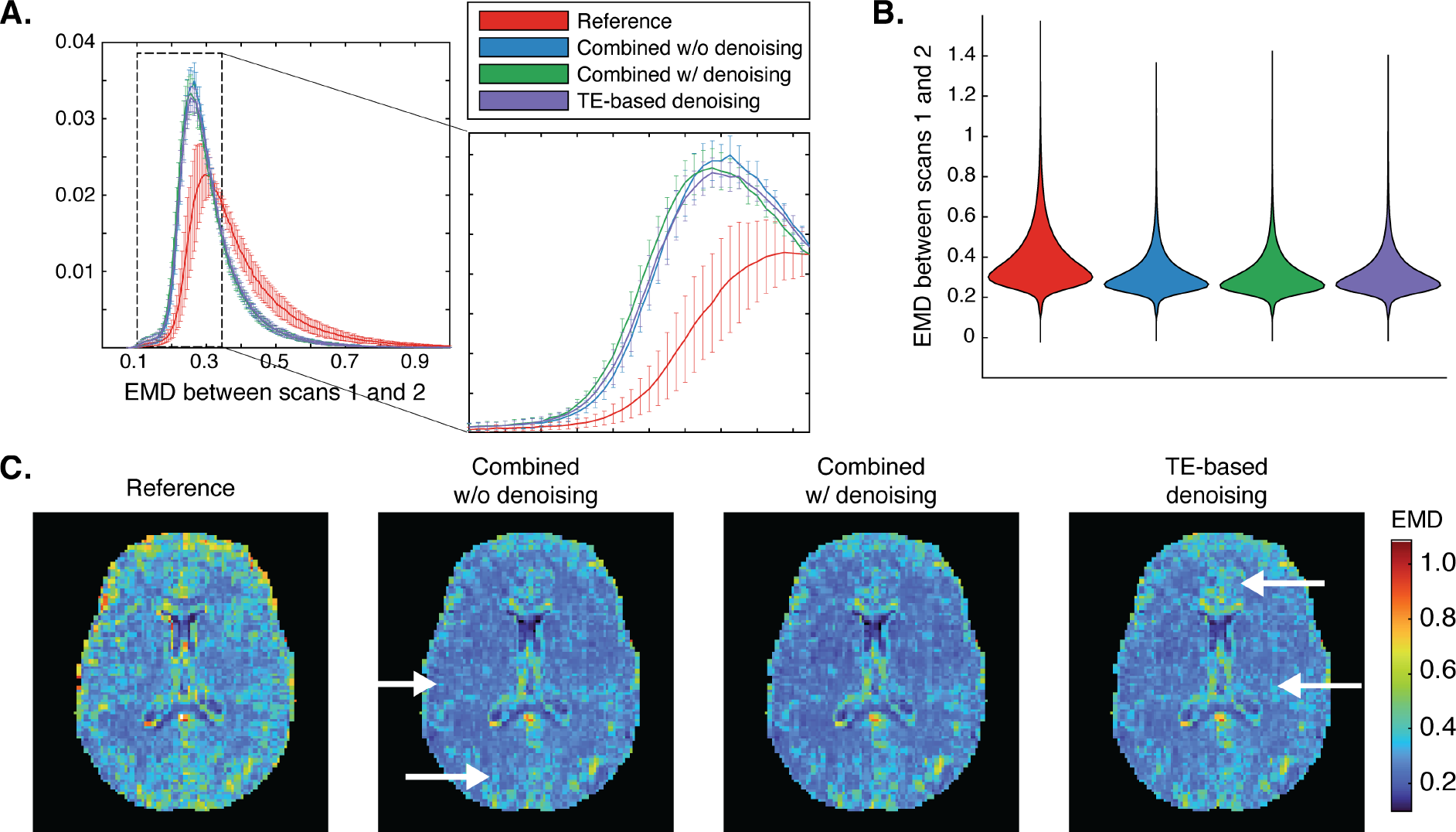
Assessing the impact different denoising strategies had on the MD-MRI pipeline and estimates. (A) Whole brain EMD histograms averaged across the five subjects for each denoising method. Error bars represent standard deviations. (B) Voxelwise EMD distributions across the study population for each denoising strategy. (C) Maps of the voxelwise EMD between **D**(*ω*)-*R*_1_-*R*_2_ distributions at *ω* = 6.6 Hz over two scans from a representative subject for each denoising strategy. High intensities correspond to low **D**(*ω*)-*R*_1_-*R*_2_ distributions reproducibility. White arrows point to the areas with elevated variability.

Voxelwise EMD distributions for each denoising strategy are presented in Fig. 6 B. One-way ANOVA showed that the source of most mean squares variability was between groups (*p* < 0.00001), with EMD means of 3.89e-1, 3.14e-1, 3.10e-1, 3.15e-1 for the Reference, combined data without and with denoising, and TE-based denoising, respectively. Pairwise comparisons using a multiple comparison test with the Bonferroni method revealed that while the EMD obtained from denoising the combined MD-MRI data was significantly lower than the other three strategies, combining the data without denoising and performing TE-based denoising did not significantly alter the EMD.

Figure 6 C displays maps of the voxelwise EMD between **D**(*ω*)-*R*_1_-*R*_2_ distributions at *ω*/2*π* = 6.6 Hz over two scans from a representative subject using each denoising strategy. High intensities correspond to low **D**(*ω*)-*R*_1_-*R*_2_ distributions reproducibility. The Reference strategy, i.e., no preprocessing at all, produced a large and heterogeneous variability as expected. Although less apparent to the naked eye, when compared to denoising the combined data strategy, skipping or performing TE-based denoising yielded elevated EMD levels (indicated by white arrows). Raw data images with representative combinations of TE, TR, b-value and b-tensor rank under the different denoising strategies are shown in Supplementary Fig. 2. These analyses support the phantom findings, confirming that the preferred denoising strategy here is to first combine the sparse MD-MRI dataset and then apply MPPCA denoising to all the volumes simultaneously; this approach was adopted for the remainder of this study.

Figure 7 illustrates axial parameter maps derived from voxelwise and bin-resolved statistical descriptors of **D**(*ω*)-*R*_1_-*R*_2_ distributions assessed at *ω*/2*π* = 6.6 Hz in a healthy brain. Despite the relatively small number of MD-MRI encoding volumes and higher resolution, the resulting maps exhibit minimal artifacts and align with findings from prior studies that employed lower-dimensional methods using denser acquisition strategies (de Almeida Martins et al. 2020; Martin et al. 2021; Reymbaut et al. 2021).

**FIGURE 7.**
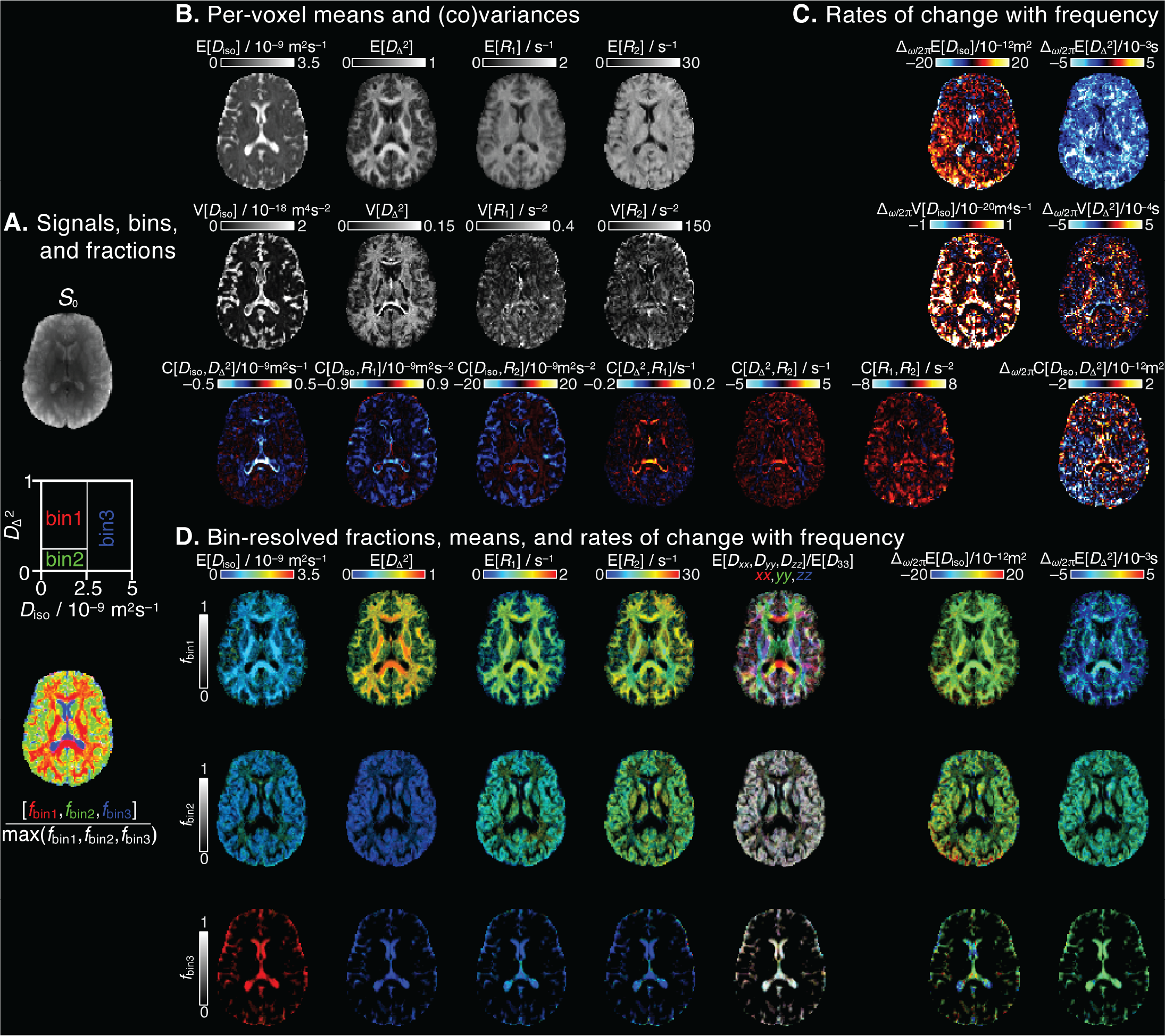
Parameter maps of a representative subject derived from voxelwise **D**(*ω*)-*R*_1_-*R*_2_ distributions. (A) *S*_0_ map displayed in gray scale, diagram with the division of the 2D 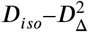 projection into three bins (bin1, bin2, bin3), and the resulting signal fractions (*f*_*bin*1_,*f*_*bin*2_,*f*_*bin*3_) coded into RGB color. (B) Per-voxel means E[x], variances V[x], and covariances C[x,y] at a selected encoding frequency *ω*/2*π* = 6.6 Hz. (C) Parameter maps of the rate of change with frequency, Δ _*ω*/2*π*_ E[x]. (D) Bin-resolved maps of E[x] and Δ_*ω*/2*π*_E[x]. The brightness and color scales represent, respectively, the signal fractions and the values of each parameter.

Intravoxel information in the human brain can be obtained and mapped by partially integrating regions of the voxelwise **D**(*ω*)-*R*_1_-*R*_2_ distributions. We chose here to be consistent with previous works and use three regions in the 2D 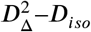 distribution space that roughly correspond to WM, GM, and CSF, referred to as bin 1, bin 2, and bin 3, respectively, and which are visualized in Fig. 7 A. When applied voxelwise, partial integration according to these bins results in their respective signal fraction maps, which are visualized as an RGB image, in which red, green and blue correspond to bin 1, 2, and 3, respectively. The RGB signal fractions map provides intravoxel information, and highlights WM, GM, and CSF contributions, as well as their superpositions.

Similar to previous studies (Martin et al. 2021; Reymbaut et al. 2021), we observed that in voxels containing multiple water populations with distinct diffusion and relaxation properties, intravoxel heterogeneity metrics V[*x*] and C[*x, y*] exhibit non-zero values. Consequently, elevated values predominantly appear at tissue interfaces, e.g., WM and CSF, with significant parameter variations that can be seen in Fig. 7 B.

The frequency-dependent findings, presented in Fig. 7 C, illustrate the rate of change with frequency within the examined range of 6.6–21 Hz for voxelwise means, variances, and covariance of *D*_*iso*_ and 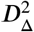. As mentioned above, posi-0078 tive Δ_*ω*/2*π*_ E[*D*_*iso*_] values indicate diffusion time dependency behavior suggestive of restriction (Aggarwal et al. 2012), while negative 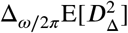 values indicate decreased anisotropy with higher frequency. We observed the highest positive values of Δ_*ω*/2*π*_E[*D*_*iso*_] in GM, predominantly in the occipital cortex. Negative Δ_*ω*/2*π*_E[*D*_*iso*_] values, indicative of incoherent CSF flow (Baron & Beaulieu 2014; Does, Parsons, & Gore 2003), were observed in the ventricles. The 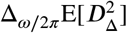 map exhibited strictly negative values, with the strongest frequency dependency in cerebral WM.

Intravoxel information can be directly imaged by resolving the diffusion and relaxation properties in Figs. 7 B and C according to the pre-defined bins. These bin-resolved means of the tensorial, relaxation, and frequency maps are presented in Fig. 7 D. This analysis parses out the characteristics of predominantly WM, GM, and CSF voxels, e.g., relaxation rates and 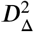 anisotropy are markedly higher in WM, compared with GM and CSF. Bin-resolved 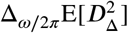 maps confirms that indeed cerebral WM regions demonstrate the strongest frequency-dependence of anisotropy.

## 4 DISCUSSION

The wealth of information encapsulated within the nonparametric **D**(*ω*)-*R*_1_-*R*_2_ distributions was demonstrated here using a diffusion phantom and healthy human subjects. Our work was focused on designing and testing a sparse and efficient MD-MRI dataset comprising 139 images, which provides full brain coverage at 2 mm isotropic voxel size, in just 40 minutes. This work presents a framework that encompasses lower-dimensional methods, i.e., 6D **D**-*R*_1_-*R*_2_ (Martin et al. 2021), 5D **D**-*R*_1_ (Reymbaut et al. 2021) and **D**-*R*_2_ (de Almeida Martins et al. 2020), 4D **D** (Topgaard 2019), and 2D distributions (English, Whittall, Joy, & Henkelman 1991; Kim et al. 2017; Pas, Komlosh, Perl, Basser, & Benjamini 2020), while including a diffusion time/frequency dependency that unlocks unique information. The efficient acquisition design prompted an inquiry into the denoising of sparse MD data, a question we have methodically explored.

Utilizing the inherent spectral content of the diffusion gradient waveforms to explore the frequency dependence of the diffusion tensor distribution and their correlations with *R*_1_ and *R*_2_, may provide crucial missing information regarding the diffusion time-dependency in biological tissue (Burcaw et al. 2015; Fieremans et al. 2016), via a unified, (biophysical) model-free framework. While conventional oscillating gradient spin echo (OGSE) protocols (Stepišnik 1981 1985) can reach a maximum frequency of 60 Hz using clinical scanners (Arbabi, Kai, Khan, & Baron 2020), the numerically optimized linear, planar, and spherical b-tensors utilized in our study, while limited to a diffusion frequency range of 6.6–21 Hz with current standard hardware, offer distinct advantages. These rank 2 and 3 b-tensors are able to encode diffusion correlations (Mitra 1995), and thus to generate the full-rank diffusion covariance tensor (Benjamini, Katz, & Nevo 2012; Ning, Szczepankiewicz, Nilsson, Rathi, & Westin 2021; Westin et al. 2014), setting them apart from OGSE-based techniques (Baron & Beaulieu 2014; Xu et al. 2016). Consequently, we have demonstrated how this frequency dependency can be dissected into intravoxel components based on their **D**(*ω*)-*R*_1_-*R*_2_ distribution, enabling the creation of microstructure-specific maps.

Although high-dimensional data is expected to contain redundancies that could be leveraged in MPPCA denoising (Olesen et al. 2023), the validity of this assertion towards the sparse MD-MRI protocol we present here had to be examined. We used the RMSE of the MD-MRI model fit from the isotropic and anisotropic ROIs within the phantom to assess the performance of several denoising strategies: denoising the diffusion data for each TE separately (Veraart et al. 2018), denoising the full MD data, and skipping the denoising step altogether. The results showed, as expected and previously demonstrated (Does et al. 2019; Schilling et al. 2023), that MPPCA denoising improves the overall performance, while indicating an advantage towards denoising the full MD data over the other strategies (Fig. 3). We used a conservative choice of the denoising window based on the original recommendation (Veraart, Novikov, et al. 2016). Such a choice, which translates to relatively small window sizes in the case of our sparse data acquisition, also avoids unwanted bias and reduced sensitivity in the output, as was recently shown (Fernandes, Olesen, Jespersen, & Shemesh 2023).

A diffusion phantom was then used to demonstrate the robustness of the sparse imaging protocol and processing pipeline, and to illustrate how the high dimensional diffusion-relaxation information can be linked to *a priori* known and well-characterized microstructure. Full distributions from single voxels in the two distinct phantom ROIs were projected onto 2D 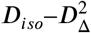, *D*_*iso*_–*R*_1_, and *D*_*iso*_–*R*_2_ planes (Fig. 4). These lower-dimensional distributions showed how the microstructural information is robustly captured via distinct spectral signatures for the isotropic and anisotropic regions of the phantom. Further, the frequency dependence of *D*_*iso*_ and 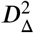 was clearly exhibited: water restricted between the fibers experience reduced interactions with barriers in the short diffusion time regime (high *ω*), resulting in increased apparent *D*_*iso*_ and in the rise of a water population with decreased anisotropy 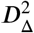. The appearance of a low diffusion anisotropy peak at high frequency provides direct evidence of the coupling of diffusion length and time scales. With assumed hexagonal close packing of the phantom’s fibers (Laun, Huff, & Stieltjes 2009), the physical characteristic length scale is about 7.6 *μ*m, while the diffusion length scales in the phantom are about 6 and 11 *μ*m for 21 and 6.6 Hz, respectively. And indeed, these numbers are supported by the relatively unrestricted water population seen at the high end of the frequency range, which vanishes as the diffusion length scale becomes greater than the physical characteristic length scale.

First and second order statistical descriptors of *D*_*iso*_, 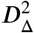, *R*_1_, and *R*_2_ were used to summarize and visualize aspects of the phantom voxelwise 6D distributions in a straightforward manner (Fig. 5). These maps were all in accordance with the expected diffusion and relaxation values (Laun, Huff, & Stieltjes 2009). The frequency-dependence map of Δ_*ω*/2*π*_E[*D*_*iso*_] is equivalent to Δ_*f*_ADC measured with OGSE (Aggarwal et al. 2012; Kershaw et al. 2013), and showed no frequency-dependence in the isotropic ROI but did show positive values in the anisotropic ROIs, as expected due to restriction. Similarly, the 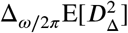 map showed no frequency-dependence in the isotropic ROI but showed negative values in the anisotropic ROIs, which is expected because the diffusion length scale at *ω*/2*π*=21 Hz is smaller than the nominal characteristic length scale of the phantom, described above.

We further assessed the performance of the investigated denoising strategies using *in vivo* human data to complement the results obtained from the phantom study. We conducted scans on five control subjects, which is close to the median sample size of six reported for technical MRI studies (Hanspach et al. 2021), each scanned twice with a few weeks in between, and quantified the variability of voxel-wise **D**(*ω*)-*R*_1_-*R*_2_ distributions between scans under different denoising strategies. The results, depicted in Fig. 6, reaffirm the conclusions drawn from the phantom-based experiments. Specifically, they highlight the preference for denoising the complete MD dataset when dealing with sparsely encoded data. While taking advantage of the inherent redundancy in each dimension of multidimensional data can enhance noise removal (Olesen et al. 2023), it necessitates dense sampling across all explored dimensions in a grid-like pattern. Our efficient MD-MRI acquisition protocol, as presented here, does not align with these conditions, as evidenced by the consistency between the phantom and *in vivo* results in favor of the same denoising strategy.

Parameter maps, presented in Fig. 7, such as E[*D*_*iso*_] and 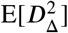 show similar trends as in previous *in vivo* human brain studies (de Almeida Martins et al. 2020; Martin et al. 2021; Reymbaut et al. 2021). As expected, bin-resolved E[*D*_*iso*_], 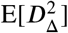, E[*R*_1_], and E[*R*_2_] all show larger values in WM (bin 1) compared with GM (bin 2). It is crucial to consider the time and length scales defined by the diffusion-encoding gradients for interpretation of the frequency dependence maps. In this work, our sampling scheme included centroid frequencies varying from 6.6 to 21 Hz (see Fig. 1), which can be used to explore restricted diffusion. The current frequency content range is expected to be particularly sensitive to length scale of approximately 13 *μ*m (if spherical geometry is assumed) (Stepišnik 1993; Woessner 1963). Therefore, the observed elevated values of Δ_*ω*/2*π*_E[*D*_*iso*_] in deep WM and cortical brain regions should be considered with respect to the probed length scale of 13 *μ*m, which would be in line with neuronal soma dimensions in the human cortex (Rajkowska, Selemon, & Goldman-Rakic 1998). Elevated Δ_*ω*/2*π*_E[*D*_*iso*_] values were also hypothesized to indicate increased local tissue disorder (Arbabi et al. 2020), which supports our findings of lower frequency dependence in WM tracts compared with subcortical WM and cortical GM. Negative Δ_*ω*/2*π*_E[*D*_*iso*_] values were observed in the ventricles, indicative of incoherent CSF flow (Baron & Beaulieu 2014; Does et al. 2003).

Basic considerations in the pulse sequence and acquisition design prevent quantitative *R*_1_ and *R*_2_ comparisons with different studies. When considering the current relaxation encoding, the measured signal arises from a complex interplay involving partial excitation by radiofrequency pulses, relaxation processes, and exchange phenomena among multiple proton pools, each possessing unique MR properties related to *R*_1_, *R*_2_, and linewidth (Manning, MacKay, & Michal 2021). These complex relationships make the measured *R*_1_ and to a lesser extent, the *R*_2_, dependent upon the particular choice of pulse sequence, slice thickness, and radiofrequency pulses bandwidth. In addition, diffusion gradient hardware limitations constrained the minimal TE in this study to 40 ms, in which myelin water (with *R*_2_ of ∼ 100 s^−1^) is expected to be fully attenuated (Manning et al. 2021). Reducing the minimal echo time can be achieved by, e.g., implementing spiral sampling of k-space (Y. Lee et al. 2021), or using next generation diffusion gradients (Huang et al. 2021).

The framework we present here is best suited when biophysical model-based approaches may not be applicable (Novikov, Kiselev, & Jespersen 2018), in which the information on the underlying tissue composition is not available beforehand. The most likely scenarios would involve pathological tissue resulting from a wide range of neurological conditions such as neurodegeneration, neuroinflammation, cancer, or even in normal aging. And indeed, diffusion-relaxation MD-MRI has been recently gaining momentum, showing that it is uniquely positioned to address a range of challenging biological questions such as prostate cancer (Wei et al. 2022; Zhang et al. 2020), breast cancer (Naranjo et al. 2021), placenta characterization (Slator et al. 2019), spinal cord injury (Benjamini et al. 2020; Kim et al. 2017), axonal injury (Benjamini et al. 2021), and astrogliosis (Benjamini, Priemer, Perl, Brody, & Basser 2022).

## 5 CONCLUSION

This study establishes the feasibility of **D**(*ω*)-*R*_1_-*R*_2_ correlation, combining diffusion-relaxation weighting, time/frequency dependence, and tensor-valued encoding, using an efficient 40-minute protocol with full brain coverage at 2 mm^3^ voxel size. The MD-MRI framework provides rich intravoxel information, without presuming tissue composition, of high-dimensional correlations of relaxation and diffusion properties, offering insights into cell types, chemical composition, axonal density, restriction, and orientations within a voxel. We demonstrated that this experimental design and acquisition protocol, in conjunction with the MD-MRI processing pipeline, produce robust estimates, encouraging the broader adoption and use of this new imaging approach in characterizing both healthy and pathological tissues.

## ACKNOWLEDGEMENTS

The authors would like to thank Mr. Phil Cholak for facilitating the MRI scans. This work was supported by the Intramural Research Programs of the National Institute on Aging and the National Institute on Drug Abuse of the National Institutes of Health.

