## Supplementary Fig for "*In vivo* disentanglement of diffusion frequency-dependence, tensor shape, and relaxation using multidimensional MRI"

---

---

### Supplementary Figures

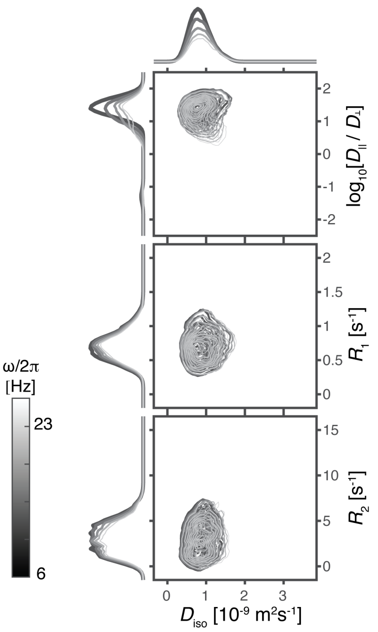


Supplementary Figure 1: Representative phantom voxel from the anisotropic ROI. $\mathbf{D}(\omega)$-$R_1$-$R_2$ distribution projected onto the 2D $D_{iso}$–$[D_{\|}/D_{\perp}]$, $D_{iso}$–$R_1$, and $D_{iso}$–$R_2$ planes for five frequencies in the range of $\omega/2\pi$= 6.6-21 Hz as indicated with the linear gray scale of the contour lines.

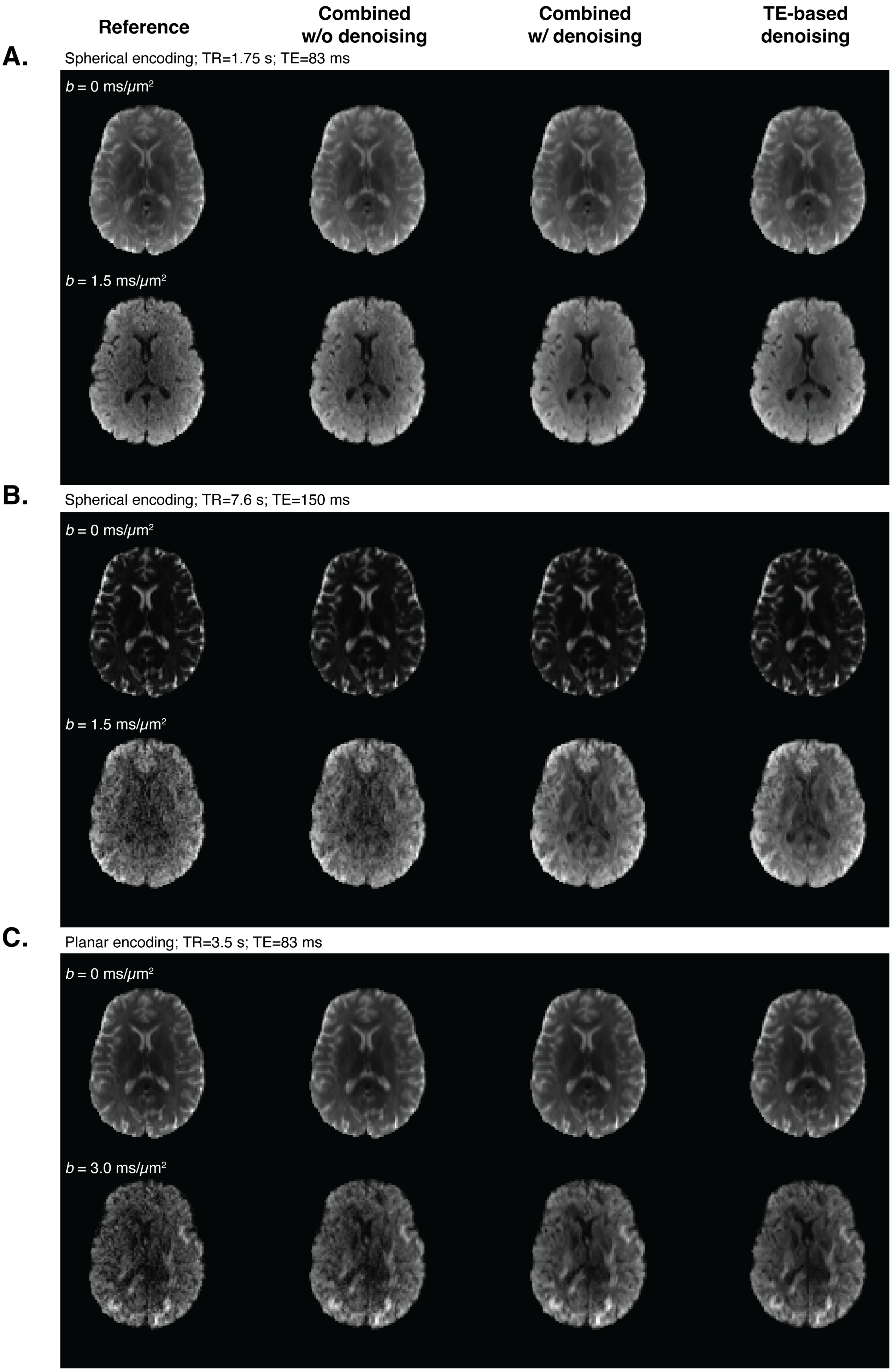


Supplementary Figure 2: Raw data images with representative combinations of TE, TR, b-value and b-tensor rank under the different denoising strategies.

**References**
